# Scalable multi whole-genome alignment using recursive exact matching

**DOI:** 10.1101/022715

**Authors:** Jasper Linthorst, Marc Hulsman, Henne Holstege, Marcel Reinders

## Abstract

The emergence of third generation sequencing technologies has brought near perfect *de-novo* genome assembly within reach. This clears the way towards reference-free detection of genomic variations.

In this paper, we introduce a novel concept for aligning whole-genomes which allows the alignment of multiple genomes. Alignments are constructed in a recursive manner, in which alignment decisions are statistically supported. Computational performance is achieved by splitting an initial indexing data structure into a multitude of smaller indices.

We show that our method can be used to detect high resolution structural variations between two human genomes, and that it can be used to obtain a high quality multiple genome alignment of at least nineteen Mycobacterium tuberculosis genomes.

An implementation of the outlined algorithm called REVEAL is available on: https://github.com/jasperlinthorst/REVEAL

## 1 Introduction

With an ever increasing length and throughput of sequencing data, affordable perfectly reconstructed complete genomes are slowly coming within reach [3, 10]. This shift in the field of comparative genomics is changing the way in which genomic variations between samples will be detected in the near future. The current approach of aligning individual short reads to a reference genome has proven useful for the detection of small variations (see Figure 1a), but will most likely be substituted by the application of *de-novo* genome assembly followed by the direct comparison of genomes. This, due to the fact that the current approach of aligning short reads to a reference genome, misses variations in repetitive or (large) structurally variant sequence (see Figure 1b). Reason for this is the fact that short reads do not span the breakpoints of large variations with respect to the reference genome. By directly comparing *de-novo* assembled genomes it is possible to detect these variations. Furthermore, by directly comparing genomes, variations can also be detected in sequence that is absent from a reference genome (see Figure 1c). This occurs when the reference genome represents an organism that is somewhat diverged from the sequenced organism of interest.

**Figure 1:**
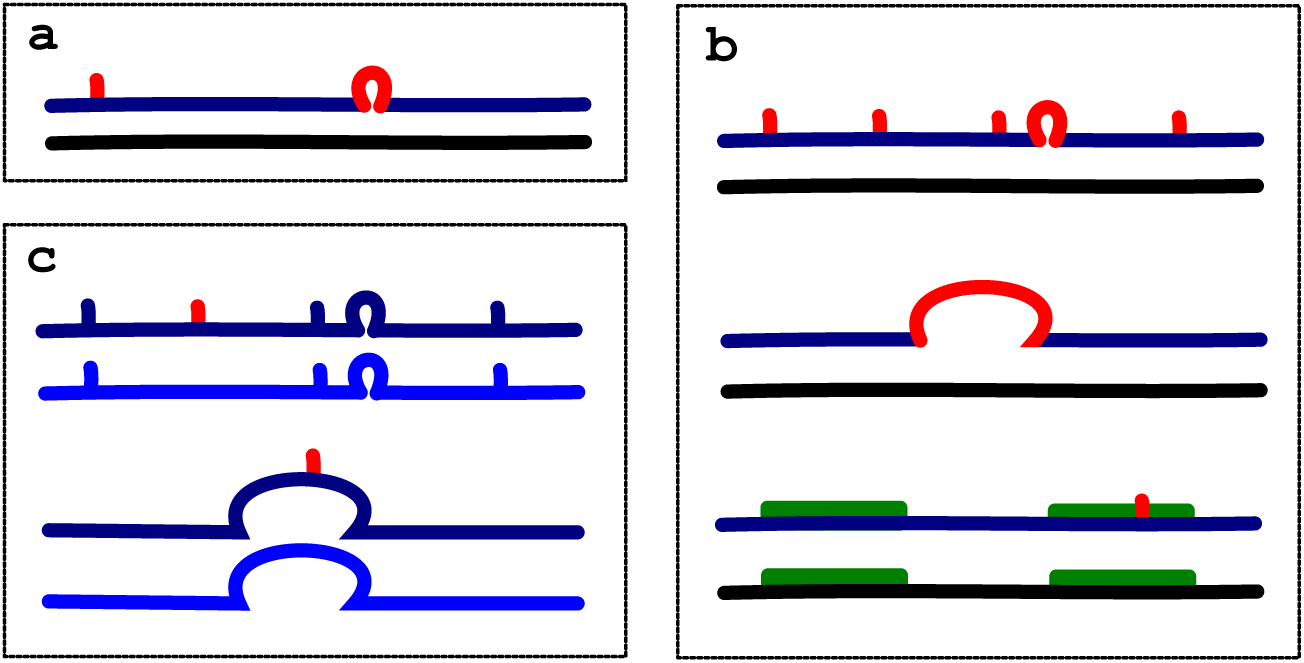
An overview of sequence variations that can and cannot be detected by the alignment of short-reads. Genomes are represented by horizontal lines, the reference genome is represented in black. Pairwise sequence variations are marked in red. **a** Short indels and point mutations can be detected by the alignment of short reads. **b** Highly variable regions, large indels and variations in repetitive regions (longer than the read length), cannot be detected by the alignment of short reads with respect to a reference sequence. **c** When comparing samples that are more closely related to eachother than to the reference sequence, simple variations between them are missed by an indirect comparison. Long reads in combination with *de-novo* assembly and whole-genome alignments allow the detection of all of these variations.

Global alignment of *de-novo* assembled genomes allows the detection of most, if not all, genomic variations. However, the computational burden of dynamic programming based global alignment algorithms like [14, 7, 12] limits its straightforward application to large eukaryotic genomes, at least when a limited amount of computational time is available. Heuristic local sequence alignment methods, like BLAST and LAST, can handle large genomes and are responsive to relatively short query sequences, but return a multitude of local sequence similarity measures, rather than the variations that differentiate between two complete genomes.

MUMmer [5], a heuristic sequence alignment method produces pairwise alignments of two entire genomes and reports variations between them. It does this by extracting all maximal exact unique matches (MUMs) between the two genomes and defines ‘chains’ (solutions of the Longest Increasing Subsequence problem) that collectively represent an alignment. Gaps between MUMs represent SNPs, indels and repetitive and/or variable stretches of sequence. This poses a problem in the alignment of genomes with high repetitiveness, and limits the resolution of variant calls. This is subsequently overcome by resorting to a dynamic programming approach for larger gaps.

Here, we introduce the concept of ‘recursive exact matching’ for global sequence alignment and provide an implementation called REVEAL. This implementation uses an hierarchical approach towards sequence alignment, which essentially builds up a tree of alignment decisions, where every knot in the tree represents an accepted MUM (see Figure 2). This way, by definition, a collinear alignment of MUMs is found without any explicit ‘chaining’.

**Figure 2:**
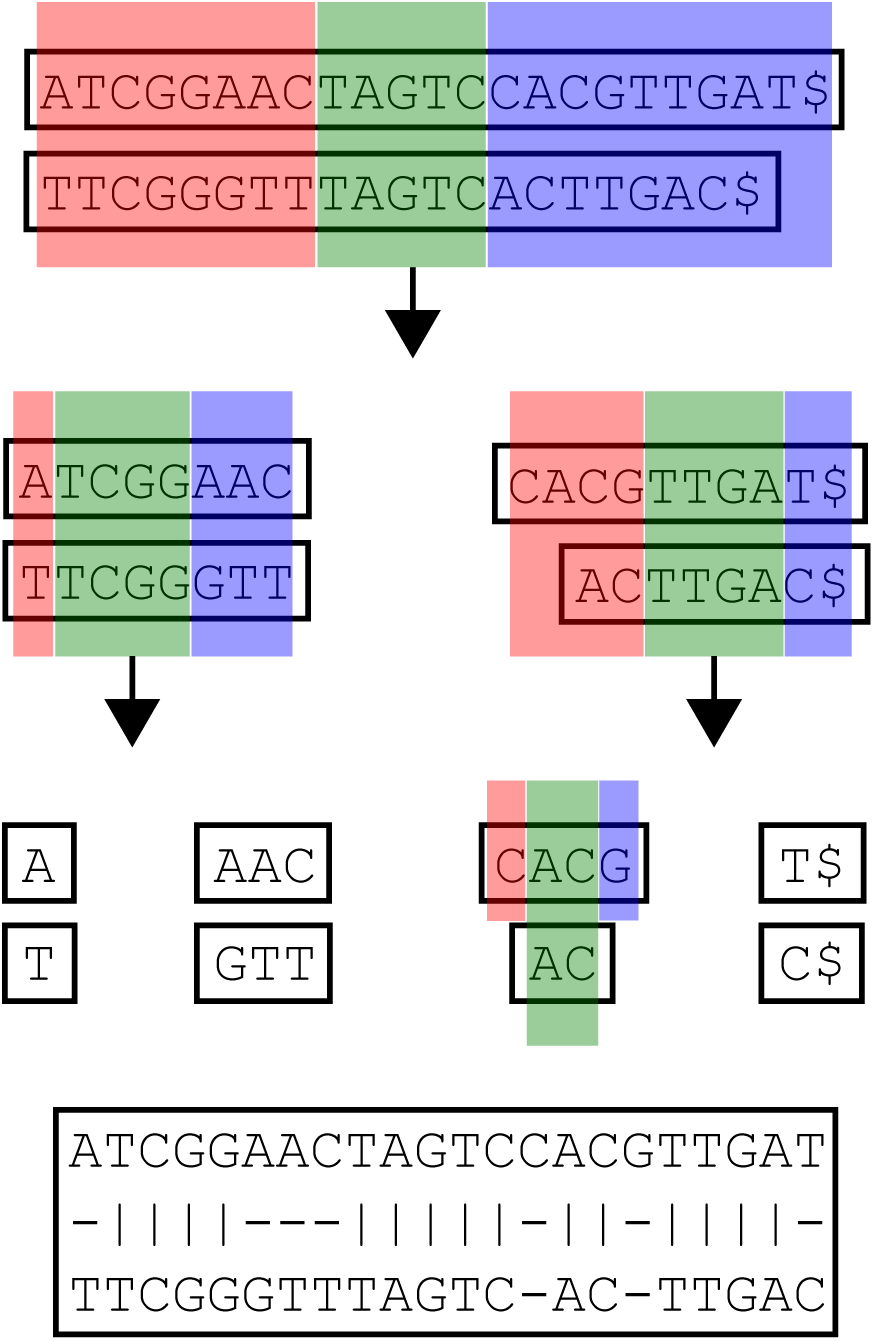
Alignment of two sequences by the application of recursive exact matching. Blue sequence is leading, red sequence is trailing, green sequence is maximally unique matching (MUM). Final alignment is obtained by four recursive alignment steps.

The advantage of this method is that repetitive stretches of sequence, that were not spanned by MUMs at the genome scale, can be spanned by MUMs in the local context. This results in higher resolution alignments without the need for dynamic programming approaches. On top of that, our method encodes the alignment in a graph structure (see Figure 3), which enables the extension of our method to construct multiple sequence alignments in a progressive way.

**Figure 3:**
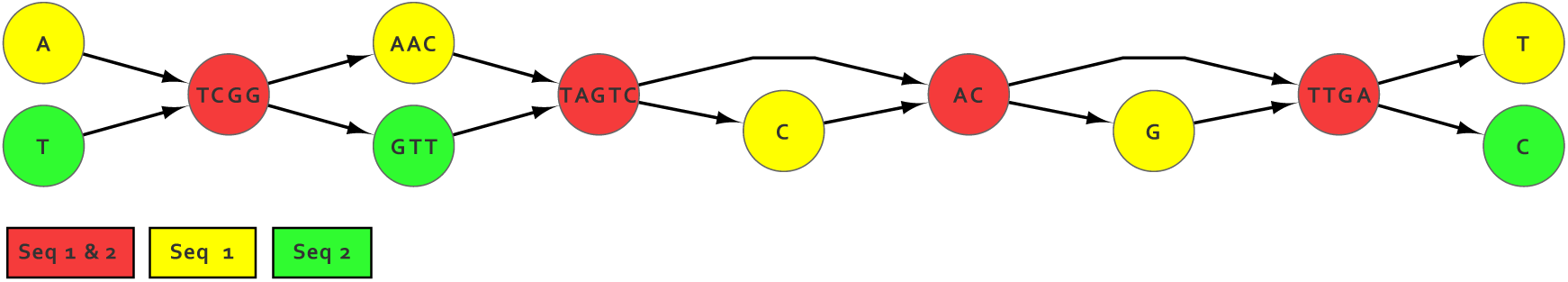
Resulting alignment graph of the two sequences mentioned in Figure 2. Green nodes are subsequences specific to input sequence 1. Blue nodes are specific to sequence 2. Red nodes are subsequences that occur in both input sequences, i.e. the hierarchically detected MUMs.

A probabilistic model based on Extreme Value Theory assesses the significance of matches throughout the construction of the alignment, and prevents the alignment of highly variable and possibly inverted sequences between the two genomes. This increases the interpretability of the alignment and aids in the translation of alignment to variant calls.

We use REVEAL to construct interpretable high-resolution alignments for human-sized genomes within reasonable time and identify many structural variations (large indels and inversions). Furthermore, we confirm the quality of our alignment using known statistics on SNPs in whole human genomes [6]. We show that REVEAL can reproduce and extend findings from a manually curated hybrid approach to detect structural variations, where individual reads originating from the same dataset were aligned and locally reassembled. Importantly, REVEAL detects many additional large inversions, and we detect a remarkable enrichment for inversions on the human X-chromosome. Finally, we show that REVEAL can also be used to construct a high quality MSA for nineteen *Mycobacterium tuberculosis* genomes, and that a MSA can be used to detect genomic variation in sequence that is absent from the H37Rv reference genome between two locally diverged samples.

Taken together, we propose a new global alignment methodology, which, through its scalability and statistical support, is especially suited for the discovery of genomic variants between whole genomes. We believe that this approach progresses the field of variant detection in third generation sequencing data.

## 2 Methods

### 2.1 Recursive MUM extraction

In order to efficiently determine the best-scoring MUM at various points of the alignment we make use of a generalized extended suffix array, which is key to keep our solution scalable. In this data structure, the indices of the suffixes originating from a concatenation of both genomic sequences (seperated by a sentinel character) are stored in a lexicographically sorted order. Additionally, a longest common prefix array (LCP array) stores the length of the common prefix between consecutive values in the suffix array. REVEAL makes use of libdivsufsort [11] to construct the suffix array and the Kasai algorithm [8] to construct an initial LCP array, however, other algorithms can also be used to generate these initial data-structures. MUMs between two input sequences can be found by a single scan over the generalized extended suffix array [1]. REVEAL essentially considers all unique exact matches within the index, scores them by their length and penalizes them by the amount of deviation from the diagonal in an imaginary alignment matrix (which essentially captures the indel size within the leading and the trailing parts of the alignment), see Figure 4.

**Figure 4:**
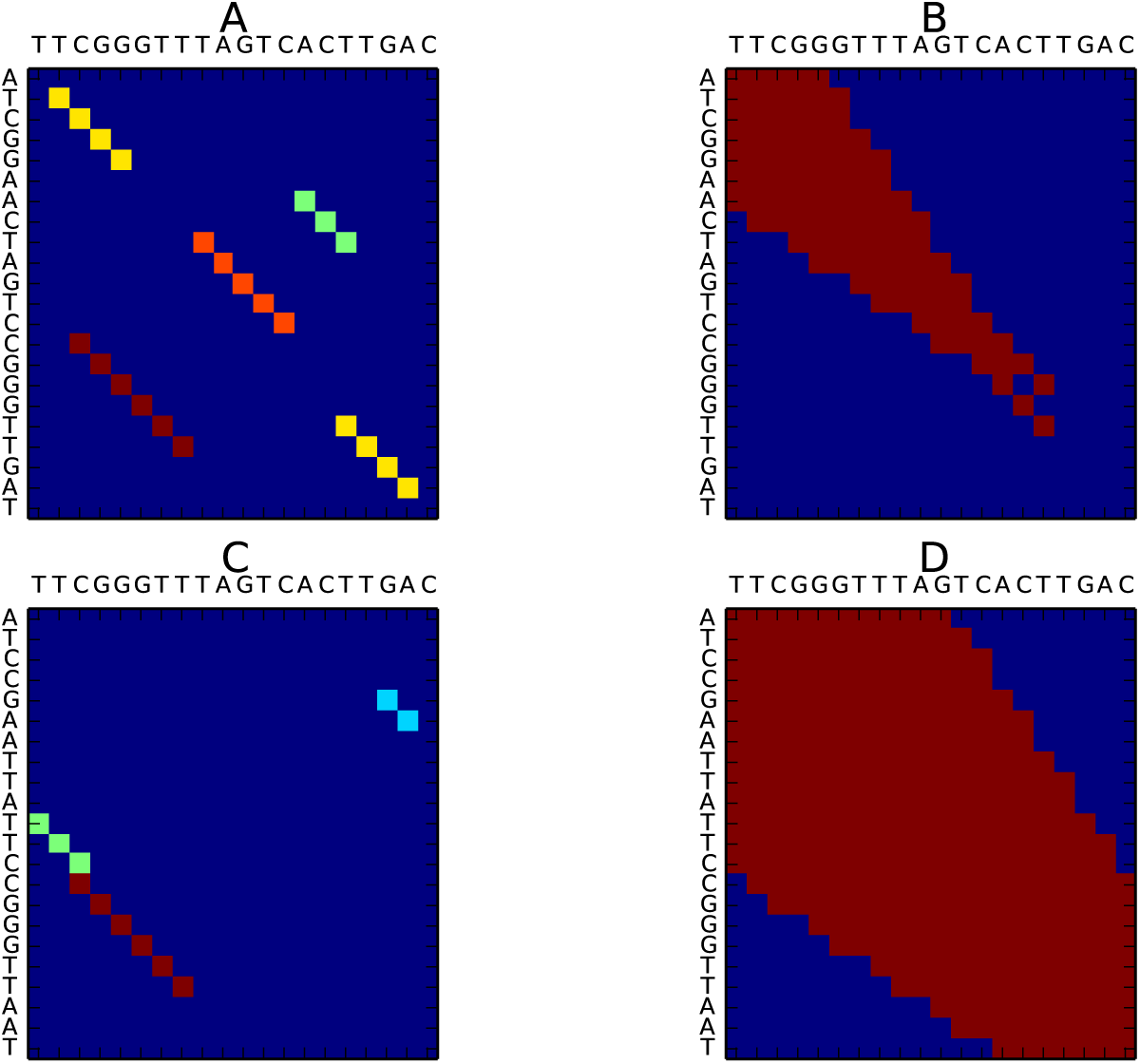
**a** The orange maximal exact match scores best when assuming an affine gap penalty model (gap open=-2 and gap extend=-1), since it is positioned on the diagonal of the matrix, while the red exact match is longer. **b** When testing the significance of the orange exact match, only the red cells of the matrix should be considered, since they are the only valid starting positions of a match that would lead to a similar gap penalty (in this example gamma=98). **c** Within this alignment the red exact match scores best (gap open=-2 and gap extend=-1). **d** When assessing the significance of the best exact match in c, more positions have to be considered as valid starting positions, because more positions within the matrix would lead to an equal or better gap penalty (gamma=318).

After the alignment of two sequences based on a single ‘best-scoring’ MUM. Data structures for the unaligned ‘leading’ and ‘trailing’ sequences can efficiently be obtained from the original extended generalized suffix array, by splitting it into two new suffix arrays for the leading and the trailing sequences (see Algorithm 1).

When we consider that the suffix array is essentially an ordered set of indices from the input text, we can split this set into suffixes originating from the unaligned ‘leading’, the aligned ‘matching’, and the unaligned ‘trailing’ sequences. Where the new LCP value can be derived from the observation that the length of the common prefix between any two values in the suffix array is equal to the minimal value on that interval within the LCP array. After splitting the suffix array, the vast majority of the suffixes will already be in the correct order. However, in some cases (e.g. with overlapping MUMs) a fraction of the suffixes has to be reordered. These can be identified, since the LCP value between two subsequent suffixes extends into the aligned part of the sequence. In these cases we use a modified bubble-sort to reorder only these suffixes and decrease their corresponding LCP values. After performing these two linear time operations we obtain two extended suffix arrays to proceed with the alignment of the unaligned leading and trailing sequences. Since the size of the suffix arrays decreases exponentially, these methods become very fast as the alignment progresses and can be performed in parallel.

### 2.2 Significance of an exact match

The alignment procedure continues until no more significant MUMs are available in any segment of the recursion tree.

At each recursion step during the alignment process, MUMs within the current sequence context are scored based on their length and penalized using an affine gap penalty model, which essentially scores MUMs by subtracting the minimal gap size created in the leading and trailing sequences (if the two sequences were aligned on this MUM) from the length of the MUM. The minimal gap size is based on the location of the MUM in an imaginary alignment matrix for the current sequence context, see Figure 4.

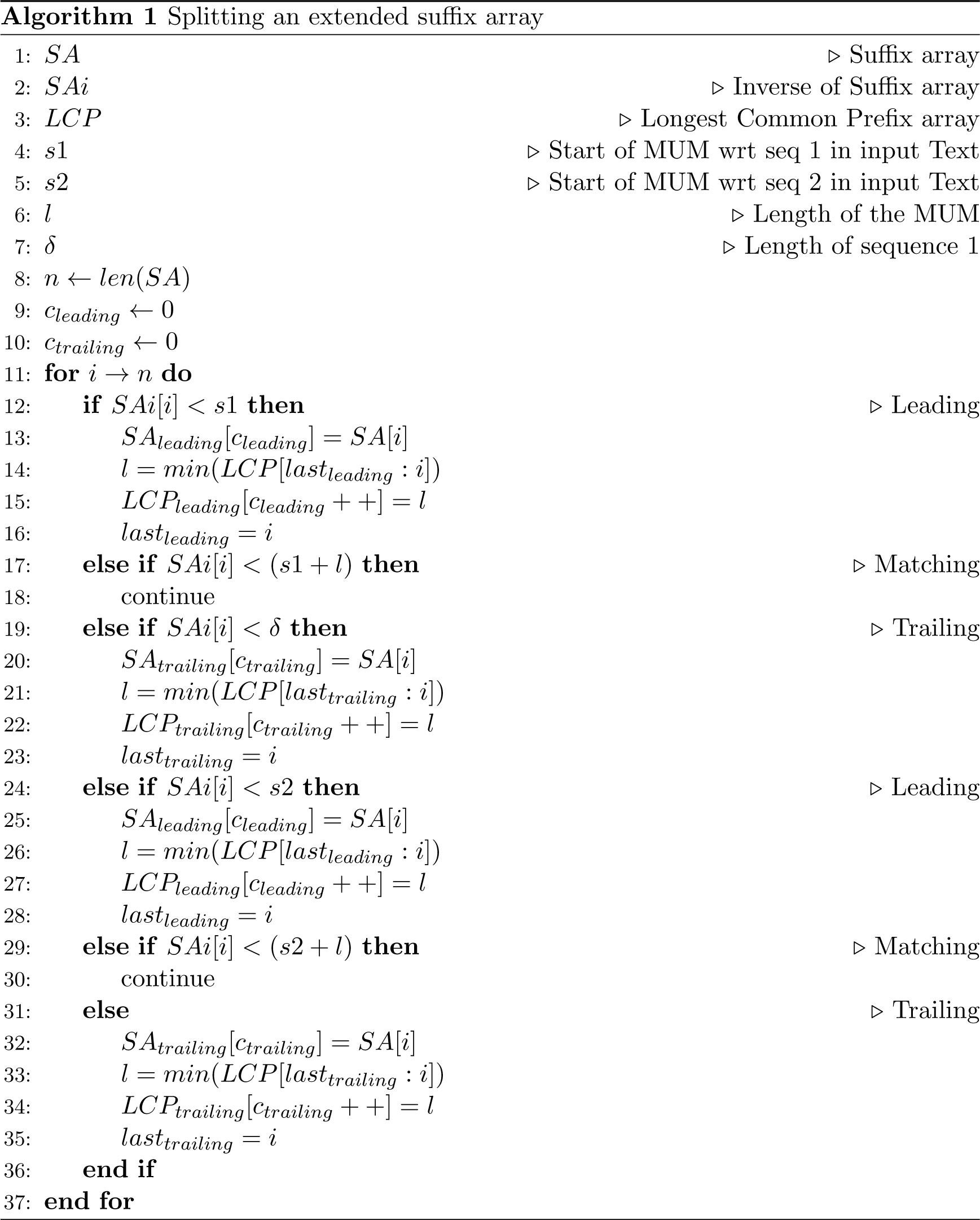

The significance of the single best-scoring MUM is then determined by calculating a p-value. This p-value is derived from a Gumbel distribution, which can be used to model the maximum expected length of an exact match between two random sequences. This allows us to determine the likelihood of a stretch of exact matching bases of a length at least as large as the best-scoring MUM. This likelihood is based upon the number of possible sites at which sequence matches of similar length as the best-scoring MUM can occur between the two aligned sequences. To incorporate the gap penalty into this model, we note that with increasing gap penalties, MUMs need to be longer to attain a similar score as the best-scoring MUM. For such longer MUMs, there are therefore less potential matching locations. Incorporating this notion in the parameters of the Gumbel distribution increases the significance of MUMs with low gap penalty values.

More specifically, the cdf of the Gumbel distribution is given by 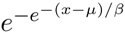 where *x* is equal to the length of the observed MUM and the parameters *μ* and *β* can be derived as follows:

*μ* = log(1*/α*) where *α* is the probability of observing a matching base between two random sequences. For now, we ignore the effect of repetitiveness and deviations in nucleotide frequency throughout the genome when modeling our null-distribution and assume a constant value of 0.25 for *α*. However, we recognize that a larger value for *α* would be more appropriate in most cases.

*β* = log(*μ*)*/γ*, where *γ* is the number of positions on which two sequences of length *m* and *n* can be compared in order to produce an exact match with a score that is equal or better than the observed match. The applied gap penalty model and the length of a MUM influence this value. The appropriate value for *γ* is best understood by considering the comparison of two sequences within a matrix, where any deviation from the diagonal can be considered as a gap in the alignment. Figure 4 shows the effect of a gap penalty model on the best-scoring MUM and the starting positions that need to be evaluated in a statistical test that determines how likely it is to observe an exact match with this length at a similar position.

Note that for off-diagonal positions only longer MUMs can attain a more significant score than the best-scoring MUM. Although it would be possible to calculate an exact p-value by using a mixture of Gumbel distributions, we approximate this value for computational reasons in a conservative manner using a single Gumbel distribution, with a cutoff corresponding to the smallest allowable MUM length.

### 2.3 Alignment graphs

The alignment of two sequences can be modeled in an alignment graph. Here we suggest an acyclic directional alignment graph in which nodes describe the sequence specifiedby an interval within one or multiple input sequences. Edges describe the contiguity of these intervals within the original input sequence.

As can be seen from Figure 3, the alignment graph compresses (in this case) two input sequences, where both input sequences correspond to two partially overlapping walks in the graph. This graph is acquired by iteratively breaking the two input nodes up into the unaligned leading and trailing sequences and collapsing the best-scoring MUM into one node, as described in Figure 2. Within the resulting graph, bubble structures indicate variations between the two sequences.

### 2.4 Alignment of graphs

Apart from the alignment of two genomes, REVEAL can also be used to align a genome to an earlier obtained alignment graph, or align two alignment graphs to eachother. This enables the construction of a multiple sequence alignments. In more or less the same way as we indexed two genomes before, we can also index the two sets of sequences corresponding to the nodes in both graphs. Next, the best-scoring MUM can be detected, followed by the alignment of the originating nodes (see Figure 5). Now a graph search (depth-first), from the aligned node is conducted in both directions in order to segment the two graphs in an ‘unaligned leading subgraph’ and an ‘unaligned trailing subgraph’. Within these subgraphs the same recursive approach can be applied until no more significant MUMs are detected. The resulting ‘multi-sample alignment graphs’ now contain all variations between all samples, which might lead to more complex and nested bubble structures that essentially describe the differences between all samples at that position, given the obtained multi-alignment. Many variations obtained in this way, are variations that cannot be detected when variants are only detected with respect to a single reference sequence (Figure 5).

**Figure 5:**
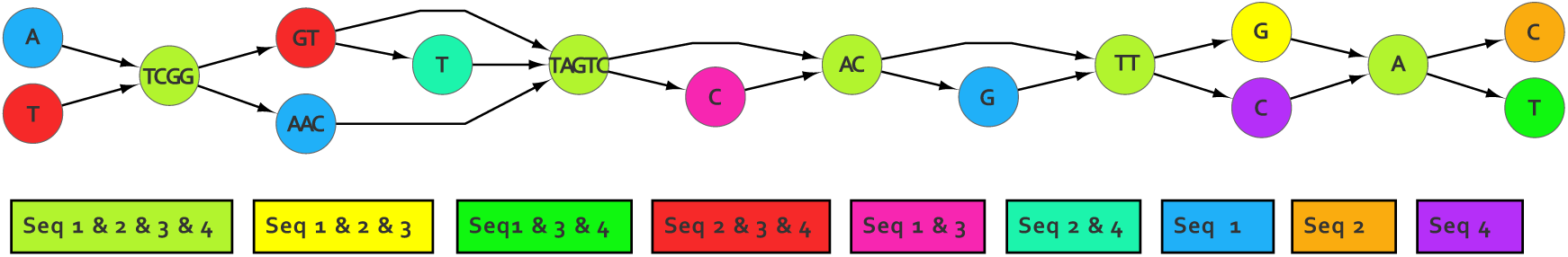
The alignment of two additional sequences (with respect to Figure 3) ‘TTCGGGTTAGTCCACTTGAT’ (sequence 3) and ‘TTCGGGTTTAGTCACTTCAT’(sequence 4) leads to a more complicated bubble structure. If we would consider input sequence 1 (‘ATCGGAACTAGTCCACGTTGAT’) as a reference sequence to which other sequences are compared one-by-one, there would not be a straightforward way of detecting the deleted ‘T’ at position 7 in sequence 3 with respect to sequences 2 and 4.

### 2.5 Alignments to variant calls

Bubbles in the resulting alignment graphs represent variations at the base pair level between genomes. Bubbles are formed by node pairs denoted as source and sink nodes. A source-sink pair is found when all paths starting in a node that is common to more than one sample, ends up in another node that is common to at least a subset of these samples. From simple bubble structures (Figure 3) it is easy to recognize SNPs and indels. Complex or nested bubbles that occur in multi sample graphs, as shown in Figure 5, can be more complicated, since they essentially describe a multitude of different pairwise variations. In the current version of this software, these bubbles are not translated into specific variant calls, but these will be addressed in future work. Simple bubble structures (also when they are nested in more complicated bubbles) are detected and translated into either substitution or indel calls between subsets of the aligned samples.

### 2.6 Detection of inversions and translocations

Substitution calls can either contain a single base (in which case they are SNPs), or contain stretches of highly variable sequence (for example, the AAC/GTT bubble in Figure 3). When these stretches are longer, they could be due to inverted sequence. In order to detect these inversions, the variant alleles in their current orientation as well as in reverse-complemented orientation (one of them) can be aligned using a simple Needleman-Wunsch alignment. If the score of the reverse complemented alignment is now considerably larger than a certain cutoff, an inversion is detected. This detection is implemented within the current variant caller (since Needleman-Wunsch alignment is used, the size of the detectable inversions is limited depending on available RAM). Translocations can also be recognized in a similar way by aligning every insert variant onto every delete variant using a sequence similarity search application like BLAST.

## 3 Results

Human-sized genome alignment possible: To show that REVEAL scales up to large sized genomes, two human genome assemblies were aligned and variations between them were analyzed. For the alignments, the manually curated human reference genome GRCh37, created by BAC cloning and Sanger sequencing, was aligned with a recent assembly of a hydatidiform mole (CHM1), using third generation sequencing [2]).

The CHM1 assembly used here consists of about 25.000 contigs. To make efficient use of the alignment algorithm proposed here, genome assemblies should be complete. Meaning that, ideally, the number of contigs resulting from a *de-novo* assembly should be equal to the number of chromosomes in the sequenced organism. To correct for this, the ordering and orientation of contigs was derived from the GRCh37 genome.

Hereto, all MUMs with a size larger than 1000 base pairs with respect to the GRCh37 genome were obtained from a suffix array that contained a concatenation of all contigs (in both orientations) and all chromosomes (in one orientation). Then, contigs were assigned to the chromosome on which most MUMs aligned. Next, contigs were ordered and oriented with respect to the reference sequence. This way, 23 chromosome-length sequences were obtained for the CHM1 assembly, which were aligned to the corresponding chromosomes on GRCh37.

For each aligned chromosome, variations were detected in the alignment graphs. From these variations, we found that the transition to transversion (Ts/Tv) ratio over all detected SNPs was 2.1 which is in accordance with Ts/Tv rates detected by other whole-genome sequencing studies of the human genome [6]. Furthermore, we observed a mutation in, on average, every 1003 bases, which is in line with an estimated average mutation load of 0.001 [6].

### Many more large indels detected

For the remaining variations we were interested in those larger than 50 base pairs. Figure 6 shows two plots of the distribution of all indel sizes larger than 50 base pairs, illustrating that more than 14000 large indels could be detected in the constructed whole-genome alignment. The distributions shown in Figure 6 reveal two peaks at around 300 and 6000 base pairs. These correspond to indels caused by ALU and L1 transposable elements. This finding was confirmed in [2].

**Figure 6:**
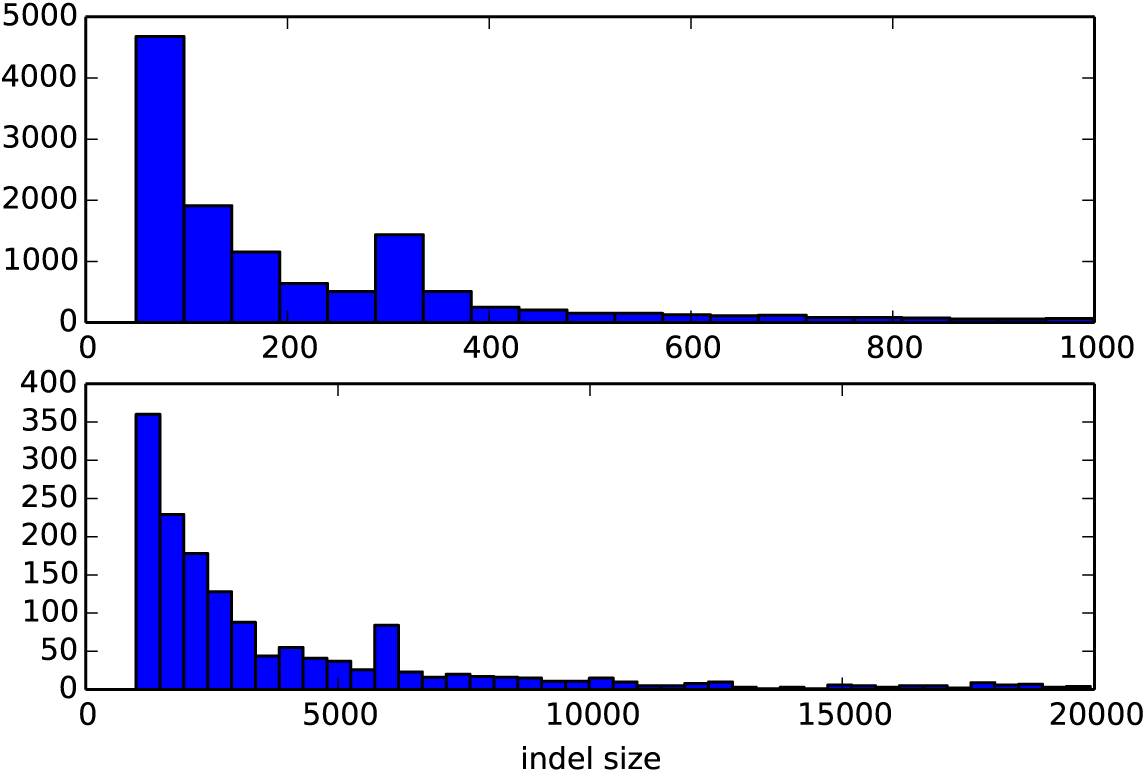
Size distributions of indels larger than 50bp between two human genome assemblies. Indels less than 1kb are shown on the top and indels greater than 1kb are shown on the bottom.

### Large inversions detected

Large inversions (up to 75.000 base pairs) were also detected. In Figure 7 we compared the number of detected inversions from the whole-genome alignment to the number of detected inversions by Chaisson et al [2]. We note that our method detects far more inversions. Remarkably, we find an enrichment for large inversions on the X-chromosome. This difference may be explained by the difference between the hybrid approach employed by Chaisson and the method used here. Since inversions are often flanked by very large inverted repeat structures, this might obfuscate the detection of breakpoints necessary for the hybrid method to detect them. Another explanation might lie in misassembly of the Celera de-novo assembler [13] that was used to construct the CHM1 genome. However, a twofold enrichment of large inversions on the X-chromosome was previously observed by a study that focussed on structural variations between human genomes of different origins [9]. In this study, this finding was explained by the enrichment of unusual inverted repeat structures on the X-chromosome, which supposedly increases the odds of structural rearrangements like inversions [16] [15]. The hybrid approach employed by Chaisson et al. might have missed this, which would make yet another strong case for whole-genome alignments of complete de-novo assemblies.

**Figure 7:**
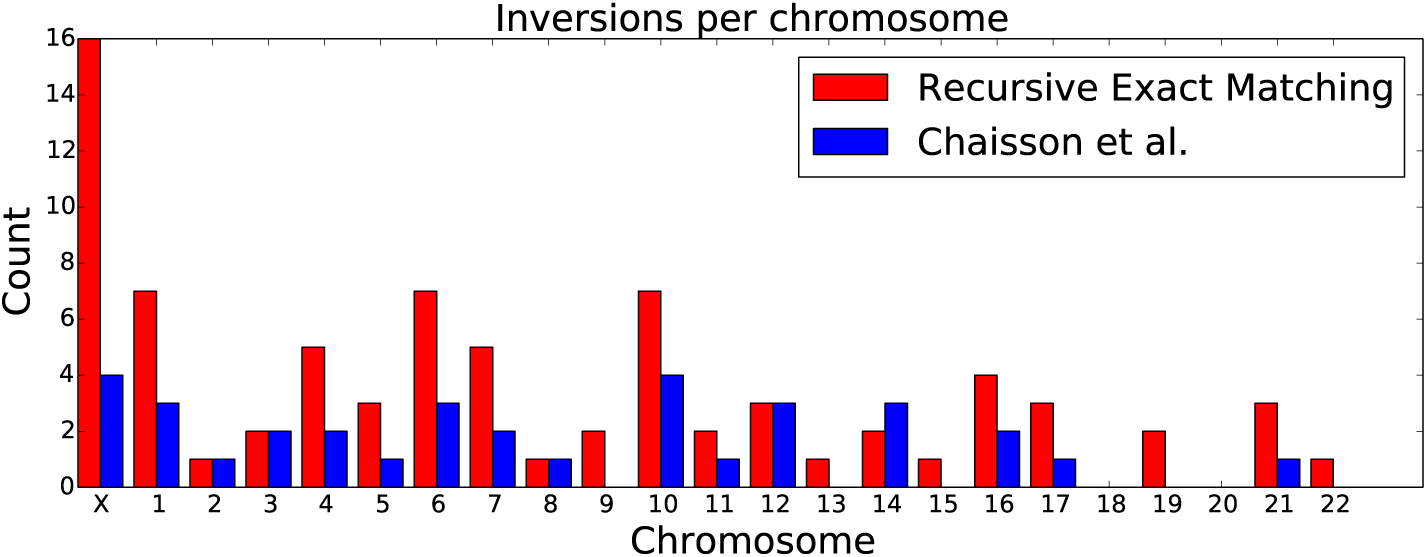
Inversions larger than 50 base pairs per chromosome. Whole-genome alignment detects more inversions than the method employed by Chaisson et al. The X-chromosome seems to be enriched for large inversion events.

### Multi alignment of 19 whole genomes possible

To show that our method does not only scale to large size genomes, but can also be used to align large numbers of genomes, we progressively aligned nineteen *Mycobacterium tuberculosis* genomes. Publicly available genomes were obtained through the TB-ARC K-RITH initiative of the Broad Institute. From these genomes, nineteen were selected from a random clade in the phylogenetic tree. Contigs were then ordered and oriented with respect to the H37Rv reference genome. Next, all genomes were sequentially aligned in a random order, resulting in an alignment graph containing a representation of all genomic variation.

### Sequence identity decreases linearly with number of genomes

By keeping track of the amount of sequence that is common to all genomes in the MSA graph we measured sequence ‘identity’ throughout the progressive alignment. As can be seen from Figure 8, there seems to be a steady linear decrease in sequence identity when adding multiple samples, as would be expected from the increase in variable sequence. Eventually, all genomes were aligned with about 93% sequence identity. We should note that the order in which genomes and graphs are aligned influences the resulting MSA, we believe that results could be further improved when this is taken into consideration.

**Figure 8:**
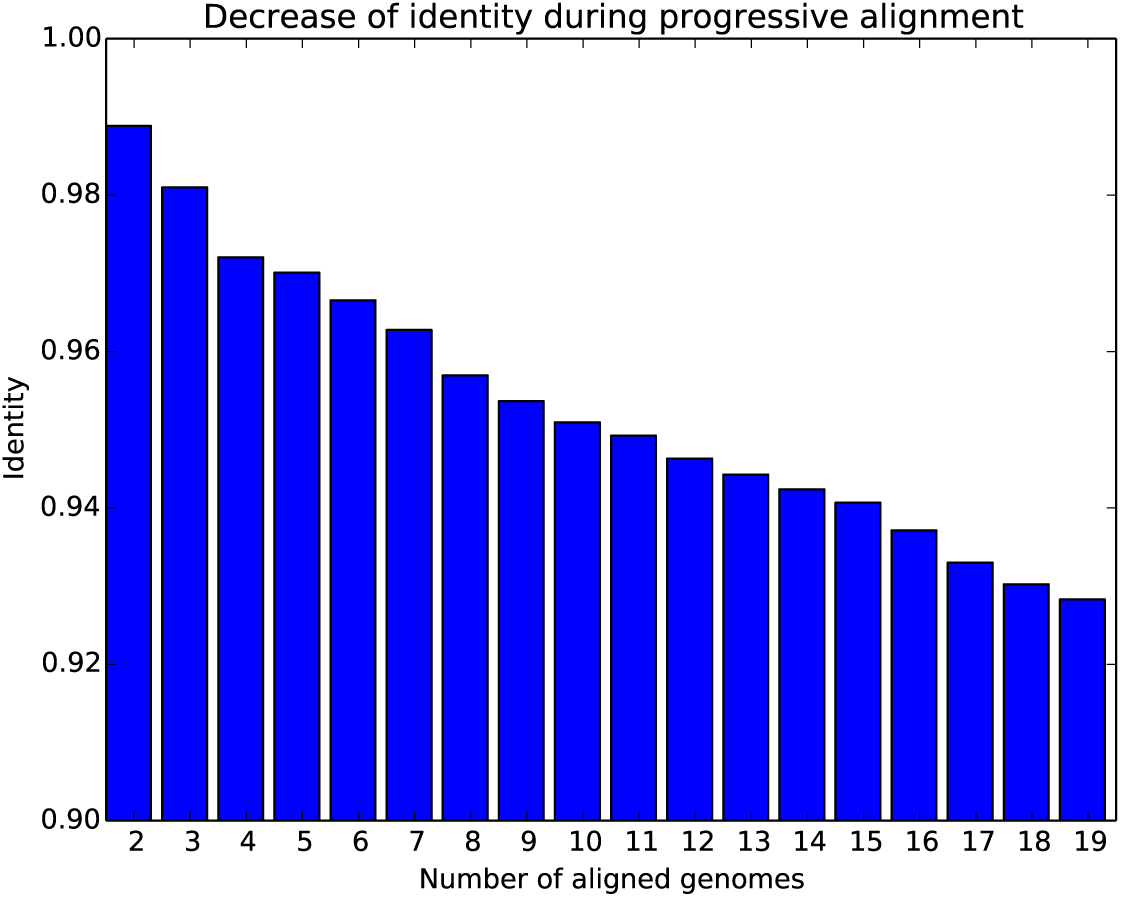
Linear decrease of identity while aligning nineteen *Mycobacterium tuberculosis* genomes

### Small sequence variation between pairs of genomes detectable

In the MSA of two different *Mycobacterium tuberculosis* genomes, *TKK 04 0085* (genbank id 622044059) and *TKK 04 0098* (genbank id 622072251), with the H37Rv reference genome, we were able to identify variations in sequence that was absent from the reference genome. In Figure 9 a bubble structure in the alignment graph shows two SNPs and one indel in sequence that is absent from the reference genome. Many more of these variations are observed in multiple sequence alignments of more distant related genomes.

**Figure 9:**
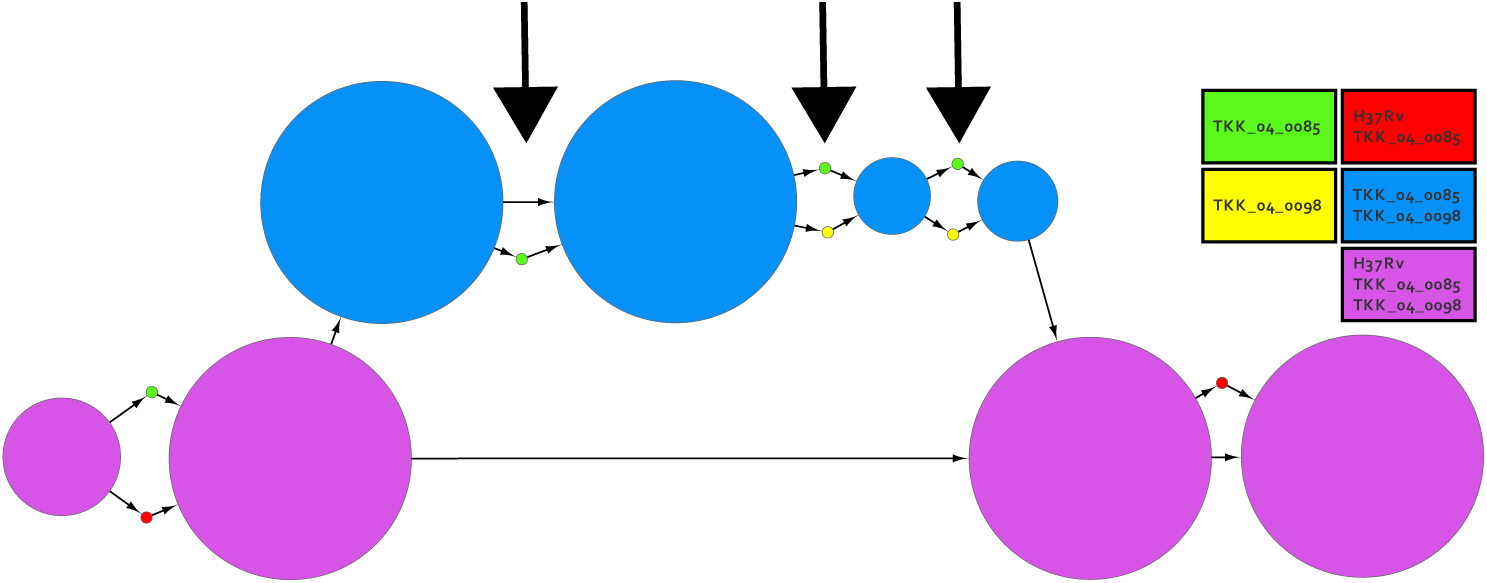
Variations between TKK 04 0085 and TKK 04 0098 in sequence absent from the H37Rv reference genome are indicated by arrows. Blue nodes contain sequence that is absent from the H37Rv reference genome but does exist in both TKK 04 0098 and TKK 04 0085. Purple nodes contain sequence that occurs in all three genomes. The size of nodes corresponds to the length of the contained sequence.

### 3.1 Performance

REVEAL uses a single CPU architecture. However, the presented algorithm is very well suited to support multiple CPU’s in parallel. All alignments presented here ran within 12 hours on a single CPU. This time can be greatly reduced by a parallel approach. The memory usage of the current implementation is about 21n bytes (O(n)), where n is the sum of the length of the two input genomes. The use of more space economical datastructures and encodings can further reduce this amount. However, we believe that the current implementation in combination with modern computer architectures will make this implementation applicable to most genomes. The sequential alignment of nineteen *Mycobacterium tuberculosis* genomes presented here took about 20 minutes on a single 2GHz Intel Core I7 processor.

## 4 Discussion

Here we introduce REVEAL, a ‘recursive exact matching’ approach that is capable of aligning whole-genomes. We show that it can be used to detect various structural variations from two *de-novo* human genome assemblies. We also showed that REVEAL can be used to align multiple genomes and illustrate the advantage of reference-free (or direct) genome comparisons by a specific multiple genome alignment of two diverged genomes.

It is important to note that, unlike exact or optimal global alignment methods, REVEAL is considered as a heuristical approach. However, this does not mean that the produced alignments are of lesser quality. In case of high sequence divergence we do indeed anticipate that an optimal alignment algorithm might be a better choice, but the optimality still depends on the minimisation of a cost function that in general only addresses substitutions and indels, but lacks inversions and translocations as biologically valid edit operations.

We believe that the concept introduced here is the way forward in order to obtain alignments of thousands to eventually millions of complete genomes that are currently being produced by initiatives like Human Longevity, Inc and others. However, we do acknowledge certain shortcomings of the current implementation.

For example, to make optimal use of the alignment algorithm, we depend on complete genomes, which means that contigs have to be ordered and oriented up front. Here, we ordered and oriented contigs with respect to an available reference sequence, however, ordering and orienting contigs with respect to another incomplete assembly, would be a more appropriate solution.

Another problem that we foresee is that we use a greedy approach towards overlapping exact matches, although the impact of this problem is limited in the case of pairwise alignments, this can become problematic in the setting of multiple sequence alignments with for example variable number tandem repeats. Multiple sequence alignments also produce more complicated (or nested) bubble structures in the alignment graph. This makes it a lot harder to make calls for all variations in a multiple sequence alignment graph.

Here, we used a progressive approach towards multiple sequence alignment. We believe that this is also the most realistic way to obtain multiple-genome alignments of thousands of large genomes. Conceptually, a recursive exact matching algorithm can also be used to align more than two genomes at a time, based on for example multi-MUMs [4]. However, this won’t scale to the number of samples that are needed to obtain statistical power in comparitive genomics studies of large genomes, and thus we believe that a progressive approach will be necessary at some point in time.

We used REVEAL to construct alignments for haplotype-genomes, the approach can also be applied to diploid genome assemblies, because they can be modeled as graphs similar to the result of the alignment of two haploid genomes. This way, alignments between a multitude of diploid genomes are conceptually possible as well.

In conclusion, we believe that REVEAL and the concepts introduced here advance the field of variant discovery from third generation sequencing data and can make an important contribution to the decoding of the second generation of complete *de-novo* assembled genomes that are currently being produced worldwide.

## Acknowledgements

We thank Jason Chin and Pacific Biosciences for providing us with the assembled contigs for the human CHM1 assembly.

